# Strength of Activation and Temporal Dynamics of BioLuminescent-Optogenetics in Response to Systemic Injections of the Luciferin

**DOI:** 10.1101/2024.04.05.588268

**Authors:** Emily F. Murphy, Aniya Means, Chen Li, Hector Baez, Manuel Gomez-Ramirez

## Abstract

BioLuminescent OptoGenetics (“BL-OG”) is a chemogenetic method that can evoke optogenetic reactions in the brain non-invasively. In BL-OG, an enzyme that catalyzes a light producing reaction (i.e., a luciferase) is tethered to an optogenetic element that is activated in response to bioluminescent light. Bioluminescence is generated by injecting a chemical substrate (*luciferin*, e.g., h-Coelenterazine; h-*CTZ*) that is catalyzed by the luciferase. By directly injecting the luciferin into the brain, we showed that bioluminescent light is proportional to spiking activity, and this relationship scales as a function of luciferin dosage. Here, we build on these previous observations by characterizing the temporal dynamics and dose response curves of BL-OG effects to intravenous (IV) injections of the luciferin. We imaged bioluminescence through a thinned skull of mice running on a wheel, while delivering h-CTZ via the tail vein with different dosage concentrations and injection rates. The data reveal a systematic relationship between strength of bioluminescence and h-CTZ dosage, with higher concentration generating stronger bioluminescence. We also found that bioluminescent activity occurs rapidly (< 60 seconds after IV injection) regardless of concentration dosage. However, as expected, the onset time of bioluminescence is delayed as the injection rate decreases. Notably, the strength and time decay of bioluminescence is invariant to the injection rate of h-CTZ. Taken together, these data show that BL-OG effects are highly consistent across injection parameters of h-CTZ, highlighting the reliability of BL-OG as a non-invasive neuromodulation method.

## Introduction

Methods of neuromodulation can provide fundamental insight into the role that neural ensembles play in mediating perceptual functions and motor actions. In a seminal study, Salzman and colleagues (Salzman et al., 1990) solidified the role of medio-temporal (MT) cortex in driving perception of visual motion by showing that intracortical microstimulation (ICMS) in MT systematically biases visual motion judgements. ICMS was also used to show that perception of flutter (i.e., low frequency vibrations) is mediated, at least in part, by neurons in primary somatosensory cortex (SI) (Romo et al., 1998). ICMS has become one of the leading methods used in brain computer interface (BCI) applications, especially in clinical applications for human patients that rely on neuroprosthetic devices (Ajiboye et al., 2017; Chaudhary et al., 2022; Collinger et al., 2014, 2013; Downey et al., 2016). Yet, although ICMS has led to fundamental discoveries, the technique can only provide uni-directional neuromodulation effects by upregulating activity of neural populations (i.e., it cannot be used to cause inhibitory effects). Further, ICMS can generate non-specific effects by modulating widespread networks through stimulation of passing fibers (Histed et al., 2009; Kumaravelu et al., 2022). ICMS also fails to provide selective modulation of cell-type specific populations (e.g., inhibitory vs. excitatory cells). Indeed, understanding how individual cell types contribute to a particular neural or perceptual function can lead to fundamental insight into the circuit mechanisms that mediate such function.

Neuromodulation methods that rely on genetic modification strategies (e.g., optogenetics and chemogenetics) can provide precise excitatory and/or inhibitory modulation of cell-type specific circuits (Aston-Jones and Deisseroth, 2013a; Bernstein et al., 2012; Bernstein and Boyden, 2011). In particular, optogenetics affords high spatio-temporal precision, even at the single cell level (Marshel et al., 2019). However, similar to ICMS, optogenetics requires an invasive surgical procedure to generate neuromodulation effects in the brain. In contrast, chemogenetic methods are minimally invasive (Berglund et al., 2016; Dimidschstein et al., 2016; Gomez-Ramirez et al., 2020; Hori et al., 2023; Song et al., 2022; Urban and Roth, 2015; Vlasov et al., 2018; Zhang et al., 2022), and provide broad coverage of activation that is particularly useful for modulating activity in large-brain animals such as non-human primates (Cushnie et al., 2023; Hori et al., 2023; Raper and Galvan, 2022). However, chemogenetics has less temporal precision, in comparison to optogenetics and ICMS, with some methods producing effects that last hours (and sometimes days) as in the case of designer receptors exclusively activated by designer drugs (DREADDs). Thus, all neuromodulation methods come with inherent advantages and disadvantages, and it is incumbent on the researcher to use the approach (or approaches) best suited to answer the question of the study.

Our recent work (Gomez-Ramirez et al., 2020) and others (Berglund et al., 2020, 2016; Crespo et al., 2021; Petersen et al., 2022; Prakash et al., 2022; Tung et al., 2015; Zhang et al., 2020) demonstrated a hybrid optogenetic and chemogenetic method that produces non-invasive optogenetic modulation using internally-generated bioluminescence. This method, termed BioLuminescent-OptoGenetics (BL-OG), tethers an enzyme (i.e., luciferase) to an opsin that is activated by bioluminescence when a luciferase catalyzes a luciferin. Coelenterazine (CTZ), the luciferin used for activating the BL-OG molecule, is innocuous and produces minimal off-target effects (Gomez-Ramirez et al., 2020). Importantly, the rapid catalytic reaction that produces bioluminescence can generate relatively short-term effects (e.g., minutes to tens of minutes). BL-OG can also provide moment-to-moment neuromodulation by conventional activation of opsins using a fiber optic cable. A critical feature of BL-OG is that the bioluminescent light created from the BL-OG mechanism provides an optical readout of the neuromodulation effect itself (Gomez-Ramirez et al., 2020). We recently showed that bioluminescent light is directly proportional to multi-unit spiking activity, and this relationship scales as a function of CTZ volume (Gomez-Ramirez et al., 2020). Taken together, BL-OG represents a highly feasible and attractive non-invasive method for generating and tracking neuromodulation effects in cell-type specific circuits.

To further establish and facilitate the use of BL-OG as a major non-invasive modulation method, it is key to establish the temporal properties and strength of the neuromodulation effects of BL-OG in response to different injection parameters of the luciferin. Here, using bioluminescence as a proxy for spiking activity (Gomez-Ramirez et al., 2020), we quantified the temporal dynamics and dose response functions to different concentrations and rate of injections of the luciferin, h-Coelenterazine (h-CTZ). Activity was imaged through the thinned-skull of a mouse to further highlight the non-invasive properties of BL-OG. We found that bioluminescence increases linearly as a function of h-CTZ dosage, and, importantly, the overall strength of bioluminescence is consistent across the rate of injection. We also observed that the onset of bioluminescence in neocortex occurs rapidly (∼10 seconds after h-CTZ injection), regardless of h-CTZ concentration. However, as expected, the onset time of bioluminescence is systematically delayed as a function of injection rate. Taken together, our data show that BL-OG effects are highly reliable across injection parameters of the luciferin, highlighting the consistency and feasibility of BL-OG as a non-invasive neuromodulation method.

## Methods

### Animals

Experiments were conducted using C57BL6J mice (N = 14) bred in the vivarium at the University of Rochester. All experimental methods are consistent with National Institutes of Health guidelines and approved by the Institutional Animal Care and Use Committee at the University of Rochester (UCAR).

### Molecular construct and surgical procedures

The luminopsin-7 (LMO7) molecule comprises a fused fluorescent and bioluminescent protein, known as NCS2 (mNeonGreen-eKL9h, a modified *Oplophorus*-based luciferase), that is tethered to the N-terminus of Volvox Channelrhodopsin 1 (VChR1; **see Figure 1A**) (Björefeldt et al., 2023). LMO7 was encoded within an adeno-associated viral 9 (AAV-9) construct under the human synapsin (h-Syn) promotor.

**Figure 1:**
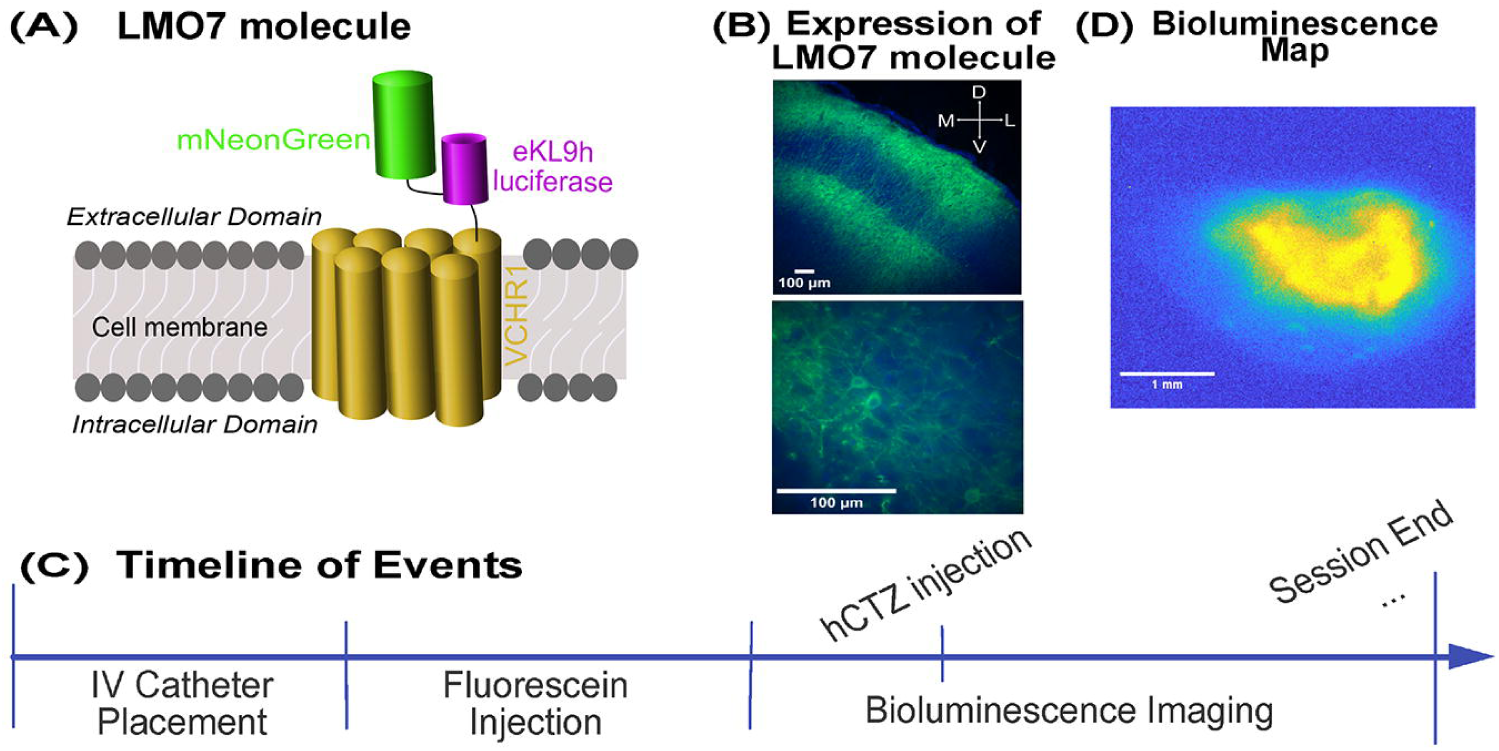
**(A)** Schematic illustration of the LMO7 molecule. The LM07 molecule comprises a fused fluorescent and bioluminescent protein, known as NCS2 (mNeonGreen-eKL9h, a modified Oplophorus-based luciferase), that is tethered to the N-terminus of Volvox Channelrhodopsin 1 (VChR1). The LMO7 molecule is anchored to the channelrhodopsin embedded in the cell membrane with the NCS2 molecule residing in extracellular space. **(B)** Representative histological images showing fluorescence expression in the left SI of a mouse. The lower panel shows a zoomed image highlighting expression of LM07 around the cell membrane. The blue oval-like shapes represent DAPI. **(C)** Illustration of the timeline of events within a single day of imaging. The session began with placing a catheter in the tail vein of the mouse, and then injecting fluorescein to verify that the catheter has accessed to the IV space. After fluorescein washed out (i.e., fluorescence was not present), bioluminescence commenced. After 10 minutes of bioluminescence imaging, we injected h-CTZ via the catheter and imaged for another 50 minutes. We continue running imaging sessions until all concentrations of h-CTZ were injected for that current day. **(D)** Representative example of a bioluminescence color map in response to a 3.93 mM concentration of h-CTZ via the tail vein.

All surgical procedures were conducted under general and local anesthesia. Mice were anesthetized with isoflurane (3% for induction and 1-2% for the remainder of the surgical procedure) and stereotaxically fixed. The fur was removed from the top of the head. The area was treated with 4 or 5% lidocaine cream, and then aseptically prepared. A custom-made head post was affixed to the skull using C&B Metabond® Quick Adhesive Cement (Parkell). The skull was shaved down until the cranium was translucent throughout a 3mm radius in the area of left primary somatosensory cortex (SI; −1.25 AP and +3.5mm ML from the Bregma). The virus with the LMO7 was injected through three burr holes in the thinned skull, 400nl in each burr hole at an injection rate of 10nl/minute. The thinned skull and burr holes were then covered with clear nail polish (Electron Microscopy Sciences, Manufacturer Part Number: 72180). Mice were imaged at a minimum of two weeks after the surgery to allow time for viral expression (see **Figure 1B**).

### Luciferin injections

Bioluminescence was produced using the substrate h-Coelenterazine-SOL (h-CTZ) for *in vivo* applications (NanoLight Technology, CAT#3011). Vials were stored at −80°C until shortly before use. Once removed from the −80°C freezer, the h-CTZ vial was wrapped in aluminum foil to protect it from light, and left on the lab counter for a few minutes to reach room temperature. One hundred microliters of sterile water was added to each vial, with vials placed in a 55°C water bath for the powder to dissolve (Crespo et al., 2021). An additional two hundred microliters of sterile saline for injection was then added to the vial to form a stock solution of 300 μL volume. For the Dose Response experiment, different levels of h-CTZ concentration (of the same volume) were derived by further diluting the stock solution with different amounts of sterile saline (e.g., 10μL and 244μL of sterile saline to achieve 3.93mM and 0.1 mM dosage concentrations of 250μL, respectively). For each concentration condition, 250μL of prepared solution was pre-loaded into a microbore tubing extension set (Smith’s Medical, 536040C), with one end connected directly to the tail vein catheter and the other end connected to another extension tubing filled with sterile water for injection. The end tube was connected to an infusion pump (Harvard Apparatus, Model 11) that controlled the rate of injection of the luciferin. For the Dose Response experiment, we injected h-CTZ with six different concentrations (0.74, 0.98, 1.72, 2.46, 3.19, and 3.93 mM). A volume of 250μL of diluted h-CTZ was injected for each dose concentration at a rate of 1ml/min. For the Injection Rate experiment, we injected two hundred and fifty microliters of h-CTZ at a concentration of 3.93 mM and rates of 0.1, 0.2, 0.3, or 1 ml/min.

### Bioluminescence and fluorescence imaging

Imaging was done through the thinned skull using an EMCCD camera (Falcon III; Raptor Photonics) fitted with a 2X objective from Thorlabs while the mouse was awake, head-fixed, and running on a 3D-printed wheel in a light-proofed chamber that was built in-house. The timeline of events is illustrated in **Figure 1C**. Prior to bioluminescence imaging, the mouse was put under light anesthesia to insert a tail vein catheter to deliver h-CTZ (Instech, Manufacturer Part Number: C10SS-MTV1417P). We injected the fluorescein dye (Ak-Fluor 10%) to confirm that the tail vein catheter was placed correctly. After observing fluorescence emitted by Ak-Fluor, the catheter was taped firmly in place, and anesthesia was removed. Bioluminescence imaging was performed approximately 20 minutes after the brief anesthesia protocol to let the animal recover and be fully awake. Unless noted, bioluminescence was imaged with 60 seconds exposure time, and with the EM-gain of the EMCCD camera set at 3000. The camera was connected to a chiller (Solid State Cooling Systems, UC160) with thermoelectric cooling temperature set to −70°C to minimize noise emitted from heat mechanisms. Micromanager was used to control the EMCCD camera (Edelstein et al., 2010). We collected ten minutes of images (i.e., 10 frames) before injecting the h-CTZ. On frame eleven, we injected h-CTZ via the infusion pump, and imaging was continued for fifty additional minutes (i.e., 50 extra frames). **Figure 1D** shows a representative example of the averaged bioluminescence map across one imaging session.

For the Dose Response experiment, we imaged bioluminescence in response to different concentrations of h-CTZ. Bioluminescence imaging in each mouse was performed across three days, with each daily experiment consisting of two or three bioluminescence imaging sessions of different h-CTZ concentrations. Daily experiments were performed approximately a week apart from each other. The highest concentration of h-CTZ was injected in each of the daily experiments. For the Injection Rate experiment, mice underwent only one experimental imaging day, with four different imaging sessions of varying injection rates (0.1, 0.2, 0.3, or 1 ml/min) using the same concentration of h-CTZ (3.93 mM) for each injection rate condition.

### Histology

Mice were euthanized with isoflurane and perfused transcardially with 4% paraformaldehyde (PFA). The brain was removed and stored in PFA at 4°C for 36 hours after perfusion. The brain was cryoprotected in 30% sucrose for another 36 hours prior to tissue slicing. Brains were sectioned in 40μm slices on a cryostat (Leica CM1850), and mounted on glass slides in mounting medium (Vectashield Vibrance with DAPI; Vector Laboratories, H-1800) and imaged using a widefield Zeiss Microscope.

### Imaging analyses

Images were taken at 16-bit resolution, and analyzed using custom-based scripts in MATLAB. For each mouse, a bioluminescence region of interest (ROI) was computed by drawing an area that contained the most amount of bioluminescence (Gomez-Ramirez et al., 2020). Note that the same ROI was used for all experimental conditions. Baseline activity was derived by averaging the signal in the ROI across the first ten frames of each experimental condition (i.e., 600 seconds). Baseline activity was then subtracted from each frame collected during the experiment.

Peak bioluminescence was calculated by averaging the image frame with maximum activity together with its two nearest neighboring frames. We performed randomization tests to statistically determine the onset and offset of bioluminescence activity in each mouse. Surrogate distributions (N = 5000) for estimating the onset of bioluminescence were generated by averaging across one hundred randomly sampled pixels from the first ten frames in the experiment (i.e., prior to h-CTZ injection). The initial frame that showed bioluminescence was estimated by determining the first two consecutive frames from the observed data that showed higher bioluminescence relative to the surrogate baseline distribution (probability value < 0.05). To estimate the offset of bioluminescence, we built surrogate distributions using image pixels that were randomly sampled from the last eight frames in the imaging session. Offset bioluminescence was estimated as the first frame of the observed data (after h-CTZ injection) that failed to show greater bioluminescence relative to the surrogate distribution built from the last eight frames in the session. Area under the curve (AUC, i.e., total bioluminescence) was calculated by averaging activity across the start and end frame of the observed data. For the highest concentration in the Dose Response experiment, the response and time dynamics of bioluminescence was estimated by averaging across the values in each of the three sessions. Statistical testing at the group level was performed using Kruskal-Wallis tests, and follow-up tests were done using Mann–Whitney U tests or curve fitting using a linear, quadratic, or exponential function. The model that provided 25% higher adjusted-R^2^ values, relative to the adjusted-R^2^ of the linear function, was determined to be the model that best explains the data. However, if none of the models provided an adjusted-R^2^ greater than 0.1, then we deemed that none of the models tested were reliable predictors of the data.

## Results

### The bioluminescence response is proportional to luciferin dosage

Bioluminescence activity systematically increases as a function of h-CTZ dosage concentration. **Figure 2A** shows the average bioluminescence time course for each h-CTZ concentration. The traces are time-locked to the onset of the injection. A Kruskall-Wallis test revealed significant differences in the bioluminescence peak response across h-CTZ conditions (H(5,47) = 22.34, *p = 4.52 x 10^-4^*; **Figure 2B**), with bioluminescence having an linear relationship to the dosage of the luciferin (adjusted-R^2^ = 0.46; **Figure 2B** *dashed lines*). We also observed significant differences between h-CTZ dosage and area under the curve (AUC, defined as the mean activity between the onset and offset bioluminescence response; H(5,47) = 31.52, *p = 7.38 x 10^-6^;* **Figure 2C**). Similar to the bioluminescence peak response, we observed a linear relationship between the AUC and h-CTZ (adjusted-R^2^ = 0.57; **Figure 2C** *dashed line*). Taken together, these data highlight the consistency and predictability of the neuromodulation effects of BL-OG in response to a particular h-CTZ concentration.

**Figure 2:**
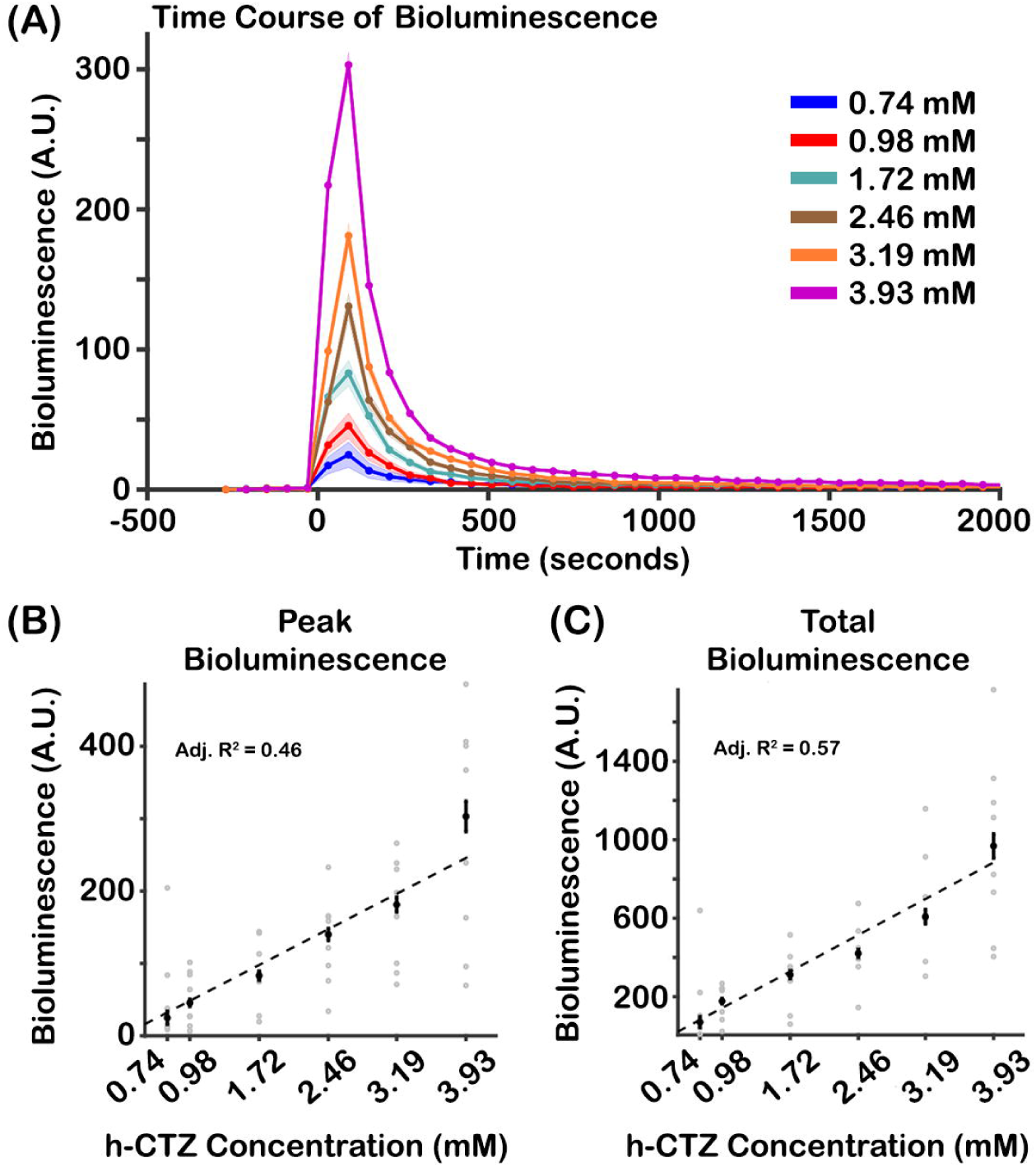
Bioluminescence response as a function of h-CTZ concentration. **(A)** Time course of Bioluminescence across h-CTZ concentration. Time point zero indicates the onset of the h-CTZ injection. Because of the course sampling of bioluminescence (i.e., 60 seconds exposure times), each dot point is plotted in between consecutive time frames. For example, the first point after the zero time point in the 3.93mM condition corresponds to 30 seconds (i.e., the middle point between 0 seconds and 60 seconds). **(B)** Average bioluminescence around the peak (± 1 frame) for each mouse across h-CTZ concentration. **(C)** Total bioluminescence response, estimated as the area under the curve between the start and end points of bioluminescent activity. The black dots indicate the median bioluminescence values across animals, whereas the gray dots indicate each individual mouse’s response. The dash lines represent the best curve fit to the data. Error bars indicate standard error of the mean (SEM). N = 8.

### Temporal dynamics of bioluminescence are consistent across luciferin dosage

Systemic h-CTZ injection generates bioluminescent activity within the first 60 seconds for all dosage concentration. **Figure 3A** shows the median onset time of bioluminescence for each h-CTZ concentration. A Kruskall-Wallis did not reveal differences in bioluminescence onset time across h-CTZ conditions (H(5,47) = 5.34, p = 0.38). Bioluminescence was imaged using a sixty seconds exposure time to obtain a strong and reliable measure of the signal (see e.g., Gomez-Ramirez et al 2020). However, this image acquisition rate provides a coarse estimate of the time onset properties of bioluminescence. As such, we performed additional imaging experiments where bioluminescence was sampled at a faster rate (exposure time = 10 seconds), at the expense of a weaker bioluminescence signal. We performed an additional control experiment where we injected fluorescein (a fluorescent dye, imaged at 0.5 Hz exposure time) via the tail vein to determine the time it takes for the injected substrate to reach the brain. **Figure 3B** shows the time course of the normalized fluorescence response (left axis, purple trace), and normalized bioluminescence response (at 10 seconds exposure, right axis, burgundy color). We observed that the onset time of bioluminescence is 10 seconds, which is largely similar to the onset of fluorescence from the fluorescein injections (∼11 seconds). Taken together, these data indicate that non-invasive injections of luciferin generate rapid BL-OG effects.

**Figure 3:**
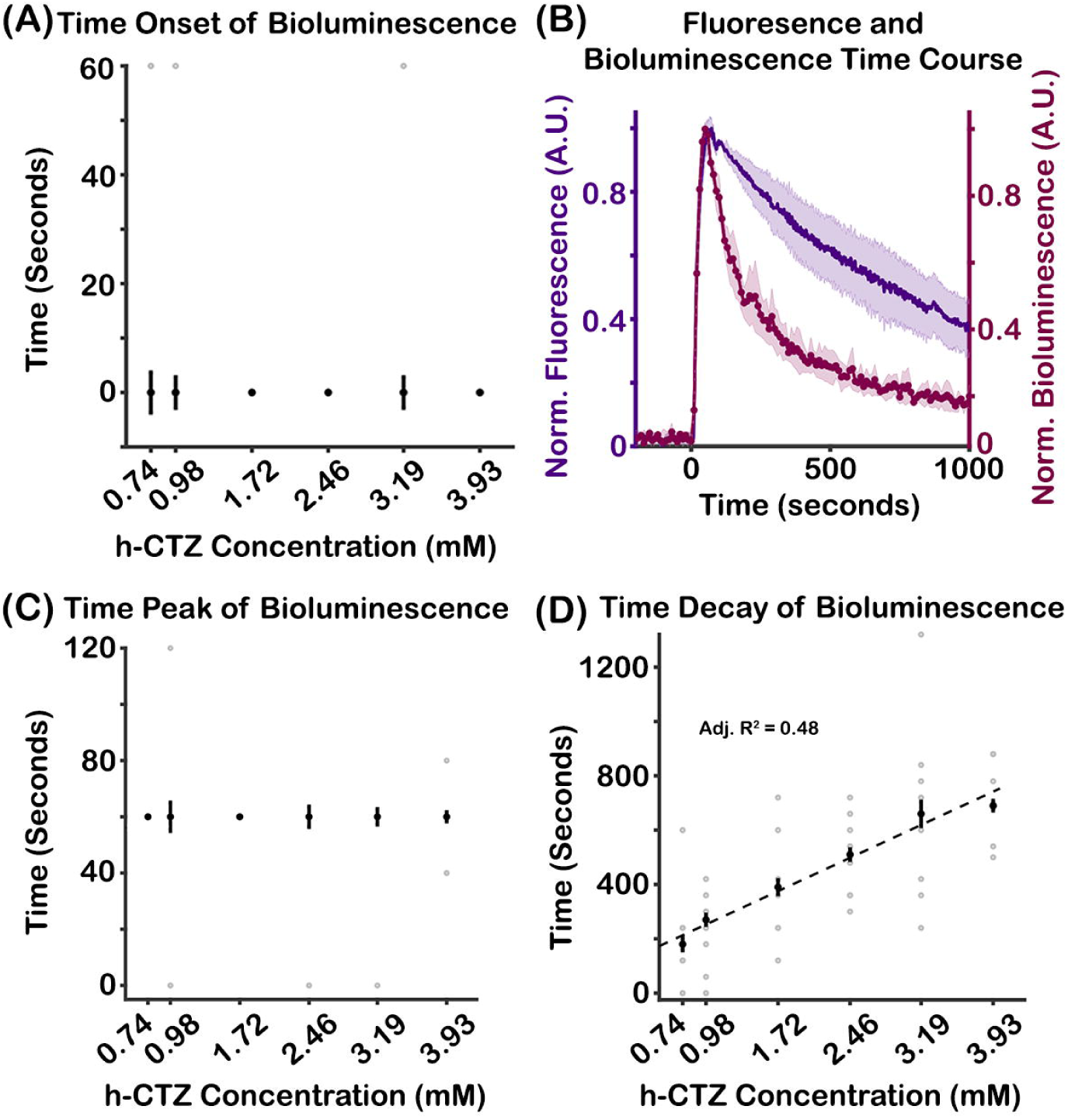
Temporal dynamics of bioluminescence response as a function of h-CTZ concentration. **(A)** Onset time of Bioluminescence across h-CTZ concentration. **(B)** Time course of fluorescence (*left axis*, from IV injection of Fluorescein) and bioluminescence (*right axis*) collected using exposure times of 0.5 and 10 seconds, respectively. The data are aligned to the onset time of the fluorescein and h-CTZ injections. **(C)** Time of the bioluminescence peak response across h-CTZ concentration. Note that two individual points in the largest h-CTZ concentration level had peak onsets at 40 seconds, a value that is inconsistent with the exposure time of the imaging experiment (i.e., 60 seconds). This value was derived by averaging across the three sessions where the largest concentration was run, in which animals had peak onsets time of 0 (N = 6 out 8), and 60 (N = 2 out 8), thus resulting in an averaged peak onset time of 40 seconds. **(D)** Time of decay of the bioluminescence response (i.e., tau) across h-CTZ concentration. Time of decay was measured as the time between the peak response and the time point where bioluminescence returned to baseline. The black dots indicate the median bioluminescence values across animals, whereas the gray dots indicate each individual mouse’s response. The dash lines represent the best curve fit to the data. Note that for Figures 3A and **3C**, the curve fitting analyses did not provide a reliable fit. Error bars indicate SEM. N = 8.

The bioluminescence peak response is also similar across h-CTZ dosage conditions (**Figure 3C**). In particular, a Kruskall-Wallis test showed no significant differences in peak activation timing across all h-CTZ dosages (H(5,47) = 2.28, p = 0.81), with the peak activity occurring between 60 and 120 seconds following h-CTZ injection. In contrast, we found that the decay time of the bioluminescence response systematically increases as a function of dosage concentration (**Figure 3D**). A Kruskall-Wallis test revealed significant differences in time decay between h-CTZ dosage (H(5,47) = 25.75, *p = 9.97 x 10^-5^*), with bioluminescence having a linear relationship to h-CTZ dosage (adjusted-R^2^ = 0.48; **Figure 3D** *dashed lined*). These data show that although the onset and peak responses of BL-OG are consistent across dosage, the duration of BL-OG effects systematically increases as a function of the luciferin concentration.

### Bioluminescence dynamics are largely stable across luciferin injections with different rates

Bioluminescence activity is similar regardless of injection rate of h-CTZ. **Figure 4A** shows the average bioluminescence time course for each h-CTZ injection rate. The traces are time-locked to the onset of the injection. A Kruskall-Wallis test did not reveal significant differences in peak responses (H(3,30) = 1.92, p = 0.59; **Figure 4B**) or AUC activity (H(3,30) = 1.56, p = 0.67; **Figure 4C**) across h-CTZ injection rate conditions. Thus, the strength of BL-OG effects appears to be insensitive to the injection rate of the luciferin.

**Figure 4:**
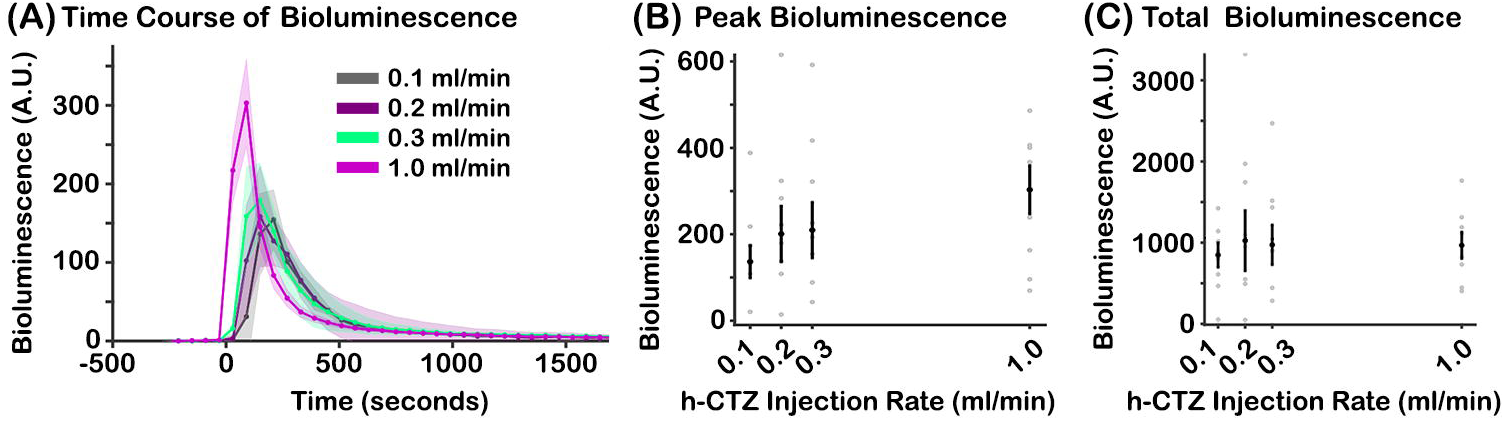
Bioluminescence response as a function of h-CTZ injection rate. **(A)** Time course of bioluminescence across rate of injection of h-CTZ (at 3.93 mM concentration). Time point zero indicates the onset of the h-CTZ injection. Similar to Figure 2A, each dot is plotted between consecutive time frames. The first point after the zero time point in the 1.0 ml/min condition corresponds to 30 seconds (i.e., the middle point between 0 seconds and 60 seconds). **(B)** Average bioluminescence response around the peak (± 1 frame) for each mouse across h-CTZ injection rate. **(C)** Total bioluminescence response, estimated as the AUC between the start and end points of bioluminescent activity. The black dots indicate the median bioluminescence values across animals, whereas the gray dots indicate each individual mouse’s response. The dash lines represent the best curve fit to the data. Note that the peak and AUC of the bioluminescence response were constant across rate conditions, thus the fits were not reliable. Error bars indicate SEM. N = 6 for injection rate conditions 0.1 – 0.3 ml/min. N = 8 for injection rate condition 1.0 ml/min.

### Temporal dynamics of bioluminescence vary by rate of luciferin injection

Injection rate of h-CTZ systematically modulates the onset of bioluminescent activity. **Figure 5A** shows the median onset time of bioluminescence for each injection rate condition. A Kruskall-Wallis revealed significant differences in bioluminescence onset time across injection rate (H(3,30) = 17.02, *p = 6.99 x 10^-4^*), with the onset time exponentially decreasing as a function of injection rate (adjusted-R^2^ = 0.51; **Figure 5A** *dashed lines*). The data further showed that the peak time of activation also decreases as a function of injection rate (H(3,30) = 17.35, *p = 5.98 x 10^-4^*, **Figure 5B**), with the bioluminescence peak time linearly decreasing with injection rate speed (adjusted-R^2^ = 0.24; **Figure 5B** *dashed lines*). However, the data did not show an effect of injection rate on the time decay of bioluminescence (H(3,30) = 3.28, *p = 0.35,* **Figure 5C**). These data reveal that the early dynamics of the bioluminescence response are systematically modulated by the rate of injection of the luciferin.

**Figure 5:**
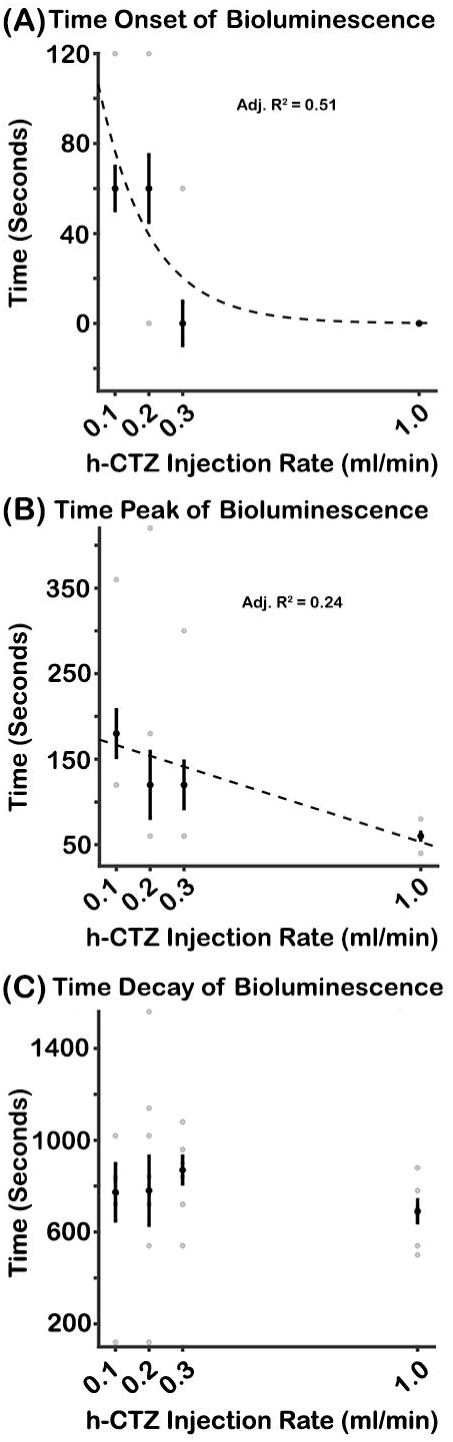
Temporal dynamics of bioluminescence response as a function of h-CTZ injection rate. **(A)** Onset time of the bioluminescence response across rate of injection of h-CTZ (at 3.93 mM concentration). **(B)** Time of the bioluminescence peak response across h-CTZ injection rate. **(C)** Time of decay of the bioluminescence response (i.e., tau) across h-CTZ injection rate. The black dots indicate the median bioluminescence values across animals, whereas the gray dots indicate each individual mouse’s response. The dash lines represent the best curve fit to the data. Note that the time of decay of the bioluminescence response was constant across rate conditions, thus the fit was not reliable. Error bars indicate SEM. N = 6 for injection rate conditions 0.1 – 0.3 ml/min. N = 8 for injection rate condition 1.0 ml/min.

## Discussion

We characterized dose response functions and temporal dynamics of bioluminescence in response to different h-CTZ injection parameters in mice expressing the BL-OG molecule LMO7, a fluorescent protein-luciferase fusion tethered to VChR1. Bioluminescence was imaged through a thinned-skull, while delivering injections of h-CTZ via the tail vein. The data show that the strength of bioluminescence, measured during the peak activity and AUC (i.e., the total bioluminescence across time), linearly increases as a function h-CTZ concentration (see **Figures 2B and 2C**). Importantly, the peak and AUC of the bioluminescence response is unaffected by the rate of injection of h-CTZ (see **Figure 4B and 4C**). Future projects using single-unit electrophysiology or 2-photon fluorescence imaging will determine whether these concentration-dependent increases in BL-OG bioluminescence lead to more neurons being recruited and/or amplifications of individuals’ neural firing themselves.

The findings in this study build on our previous work that show robust and systematic bioluminescence activity in response to direct injection of the luciferin in the brain (Gomez-Ramirez et al., 2020). Here, we further demonstrate this robust and parametric relationship between strength of the bioluminescence and luciferin dosage using systemic injections of the luciferin. Further, our data reveal that bioluminescence activity produced by the BL-OG molecule can be reliably imaged through a thinned skull of a mouse, highlighting BL-OG’s feasibility to track neuromodulation effects through minimally invasive procedures. In sum, our study provides strong evidence that IV injections of h-CTZ generate concentration-dependent BL-OG effects in the brain, and that these effects are independent of the rate at which the luciferin is delivered.

### Temporal dynamics of the bioluminescence response

The onset time of bioluminescence in neocortex is rapid and invariant to h-CTZ concentration (see **Figure 3A**). In particular, bioluminescence emerges within the first 60 seconds after h-CTZ injection in all concentration conditions. However, using shorter exposure times (i.e., higher sampling rates), we found that the onset time of bioluminescence is ∼10 seconds after h-CTZ injection. This rapid onset time highlights the robust bioavailability of h-CTZ to the brain, and the fast-acting mechanisms of BL-OG to elicit neuromodulation effects. We also found that the onset peak response is unaffected by the concentration of h-CTZ. Further, as expected, the decay time of bioluminescence linearly increases as a function of h-CTZ concentration, likely due to the large substrate amount that is catalyzed by the luciferase.

As expected, the onset time of bioluminescence modulates as a function of injection rate. Specifically, we observed a nonlinear relationship (exponential) between bioluminescence onset and injection rate, with faster rates generating earlier onset times (see **Figure 5A**). Similarly, we found that the peak time of bioluminescence decreases as the rate of injection is increased (see **Figure 5B**). However, the decay time of bioluminescence is unaffected by injection rate. Thus, our data indicate that the rate of injection of h-CTZ modulates the bioluminescence response by shifting the onset and peak activation time, without modifying the decay dynamics of the bioluminescence response. These effects of injection rate are notable because they generate bioluminescence activations (e.g., AUC responses) that are largely homogenous regardless of the injection rate (see e.g., **Figure 4C**).

The onset time of BL-OG effects appear to be substantially faster than the neuromodulation onset times of other chemogenetic methods such as DREADDs (Deffains et al., 2021; Nagai et al., 2020). For instance, it was shown that deschloroclozapine (DCZ) administered IV at a rate of 0.2 ml/sec leads to neuromodulation effects that commence around 5 minutes after injection (Nagai et al., 2020). In contrast, using the same injection rate of h-CTZ, we observed bioluminescence occurring between 60 and 120 seconds after injection (see **Figure 5A**). The difference in neuromodulation onset times between the two methods is unclear, but it should be noted that DREADDs operate on secondary messenger systems (i.e., G protein-coupled receptors; GPCRs) that have significantly slower kinetics as compared to ionotropic channels in which optogenetic elements operate on (Armbruster et al., 2007; Whissell et al., 2016). Our data also show that BL-OG effects can be short-lasting (∼3 to ∼12 minutes), providing repeated BL-OG-mediated neuromodulation effects within a single experimental day. In contrast, the duration of the neuromodulation effects of DREADDs are thought to last hours (Alexander et al., 2009; Whissell et al., 2016). Our data demonstrate that the duration of BL-OG effects can be regulated by the concentration amount, volume (see (Gomez-Ramirez et al., 2020)), or injection rate of the luciferin, indicating that the dynamics of neuromodulation effects of BL-OG can be systemically controlled by experimenters and, potentially, clinicians.

In sum, our study provides a generalized framework of the experimental parameters that control the strength and dynamics of BL-OG effects in the brain. Our data provides evidence that BL-OG effects can be fast, with the strength and duration of the neuromodulation effects modulated by the injection parameters of the luciferin. We show that BL-OG is unique in providing online tracking of the neuromodulation effects via the bioluminescent activity, and this tracking can be achieved using minimally invasive procedures. Our study highlights the reliability of BL-OG as a major neuromodulation method and demonstrates its feasibility to monitor neuromodulation effects non-invasively.

## Acknowledgements

We would like to acknowledge Drs. Ute Hochgeshwender (Professor at Central Michigan University) and Chris Moore (Professor at Brown University) for their insight in data analyses and feedback on the manuscript. We will also like to acknowledge Dr. Kuan H. Wang (Professor at the University of Rochester Medical Center) for his comments on the manuscript. This work was supported with Startup funds from the University of Rochester (MGR), and funds from the Alfred P. Sloan Foundation (MGR).

## Author Contributions

**Emily F. Murphy:** Conceptualization, Methodology, Investigation, Formal analysis, Data Curation and Writing. **Aniya Means:** Investigation and Formal analysis. **Chen Li:** Investigation and Formal analysis. **Hector Baez:** Software and Investigation. **Manuel Gomez-Ramirez:** Conceptualization, Methodology, Investigation, Formal analysis, Data Curation, and Writing

## Data and code availability

All data reported in this paper are stored in Neurodata Without Borders (NWB) format, and will be shared by the corresponding author upon request. Any additional information required to reanalyze the data reported in this paper is available from the corresponding author upon request. It is expected that publications produced from a shared data agreement would result in co-authorships for all of the authors in this manuscript.

## Declaration of Interest

None

## Notes

### Competing Interest Statement

The authors have declared no competing interest.

